# Fine-scale diversity of microbial communities due to satellite niches in boom-and-bust environments

**DOI:** 10.1101/2022.05.26.493560

**Authors:** Zihan Wang, Yulia Fridman, Sergei Maslov, Akshit Goyal

## Abstract

Recent observations have revealed that closely related strains of the same microbial species can stably coexist in natural and laboratory settings subject to boom-and-bust dynamics and serial dilutions, respectively. However, the possible mechanisms enabling the coexistence of only a handful of strains, but not more, have thus far remained unknown. Here, using a consumer-resource model of microbial ecosystems, we propose that by differentiating on Monod parameters characterizing microbial growth rates in high and low nutrient conditions, strains can coexist in patterns similar to those observed. In our model, boom-and-bust environments create satellite niches due to resource concentrations varying in time. These satellite niches can be occupied by closely related strains, thereby enabling their coexistence. We demonstrate that this result is valid even in complex environments consisting of multiple resources and species. In these complex communities, each species partitions resources differently and creates separate sets of satellite niches for their own strains. While there is no theoretical limit to the number of coexisting strains, in our simulations, we always find between 1 and 3 strains coexisting, consistent with experiments and observations.

**Author summary:** Recent genomic data has exposed to us the remarkable spectrum of microbial diversity in natural communities surrounding us, which harbor not only hundreds of species, but also a handful of closely related strains within each of those species (termed “oligo-colonization”). While the mechanisms behind species coexistence are much better studied, the mechanisms behind the coexistence of closely related strains have remained understudied. Here, using a simple consumer-resource model, we show that if strains differ on their Monod growth parameters, they can coexist even on a single limiting resource, provided that the environments vary with time in boom-and-bust cycles. The Monod growth parameters describe how a strain’s growth rate changes with resource concentration, namely the half-maximal concentration and maximal growth rate. Simulations of our model show that both in simple and complex environments, even though an arbitrary number of strains can coexist, typically it is between 1 and 3 strains of a species that coexist over several randomly assembled communities, consistent with some experimental observations. This is because the allowed parameter space for coexistence shrinks significantly with the number of strains that coexist.

## Introduction

Microbial communities in almost all natural settings are characterized by an astonishing diversity, manifesting itself at multiple evolutionary scales [1, 2]. These scales range from separate kingdoms (e.g., archaea and bacteria) all the way down to closely-related strains of the same species [3, 4, 5, 6, 7], other environments. Remarkably, this wide-ranging diversity can persist even in well-controlled laboratory settings, containing alternating cycles of exponential growth, followed by a transfer to fresh media after dilution by a large factor [8, 9]. The coexistence of distantly related community members, such as different species or kingdoms, can be readily explained via niche theory, which suggests that each species can occupy a different niche, e.g., by specializing on a different resource, allowing everyone to coexist [10, 11, 12, 13, 14]. Remarkably, it is the coexistence of fine-scale diversity — that is, closely related strains of the same species — that remains puzzling [7, 15, 16]. This is because presumably, such strains haven’t diverged sufficiently to establish and occupy distinct resource niches. Thus, the observation of fine-scale diversity suggests that every resource niche might contain several “satellite” niches, ready to be occupied by closely related strains, enabling their coexistence.

Satellite niches are not expected in stable environments, where resources are supplied continuously at a constant rate, such as in a chemostat. In these conditions, competitive exclusion guarantees that only one strain will be able to survive in every resource niche. This strain will drive down the resource to the lowest concentration, at which all other strains cannot survive [13]. In contrast, in fluctuating environments, the existence of satellite niches remains a possibility [17, 18, 19]. Indeed, in the extreme case of environmental fluctuations, i.e., in boom-and-bust cycles, the selection criteria for strains occur along two separate dimensions: one for strains that grow well at high nutrient concentrations (the “boom” phase), but poorly at low nutrient concentrations, and the other for strains that dominate in the “bust” phase, at much lower nutrient concentrations. Furthermore, strains of the same species may rapidly (with just a few mutations [20]) modify the range of nutrient concentrations optimal for their growth, allowing for rapid colonization of the available satellite niches by closely related strains, rather than by members from distant species. In conclusion, satellite niches may arise as a consequence of boom-and-bust cycles, and then be occupied by different strains of the same species, allowing their coexistence.

Here, we use a consumer-resource model of microbial communities to demonstrate how satellite niches appear and are colonized by different strains of the same species, over the process of community assembly. We start by considering the case of a single resource, and show that in this case, anywhere between 1 and 3 strains typically coexist. Indeed, the coexistence of even more strains is possible, but has a small likelihood. We then proceed to generalize our results to multiple resources (or niches), colonized by distantly-related species. In this case, each of the species comprises a small number of closely-related strains, which can coexist due to the presence of satellite niches. Our results provide a possible mechanism for the widespread observation of fine-scale diversity in natural as well as laboratory microbial communities.

## Model and Results

### Strains of the same species can coexist on a single limiting resource

To study whether multiple strains of a species — which differed in their maximal growth rate (*g*) and substrate affinity (*K*) — could coexist in boom-and-bust environments, we simulated a simple consumer-resource model inspired by serial dilution experiments. Briefly, both strains were initially inoculated in a medium consisting of one resource (e.g., carbon source) provided at a concentration *c*_0_, allowed to grow until the resource was fully depleted, and then diluted by a factor *D* and transferred to a fresh medium with resource at the same concentration *c*_0_ (Fig. 1b). We repeated these growth-dilution cycles until the community reached a steady state, that is, displaying reproducible dynamics in each cycle. Each strain with abundance *N_i_* grew exponentially, with its instantaneous growth rate determined by its Monod kinetics, its dynamics given as follows:

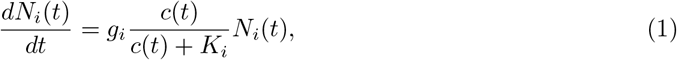

where *c*(*t*) represented the resource concentration at time *t* after the start of each cycle (*c*(*t* = 0) = *c*_0_). Here, *g_i_* represented the maximal per capita growth rate of each strain, realized at high resource concentrations (*c*(*t*) ≫ *K_i_*), and *K_i_* represented the resource concentration at which the growth rate of the strain dropped to half its maximum value. Note that the strain with a higher *g_i_* exhibits a fast initial growth, whereas a strain with a lower *K_i_* continues to grow at appreciable growth rates even at relatively low resource concentrations. Similar to previous results [21], we first considered a model with two strains of the same species, but to build on previous studies, we will later generalize this model to include multiple strains and species. In the two strain case we considered first, one of the strains (*B* in red) had a higher maximal growth rate *g_i_*, analogous to an *r*-strategist in classical ecology [22, 23], whereas the other (*A* in blue) had a much lower value of *K_i_*, analogous to a *K*-strategist in classical ecology (Fig. 1b).

**Figure 1:**
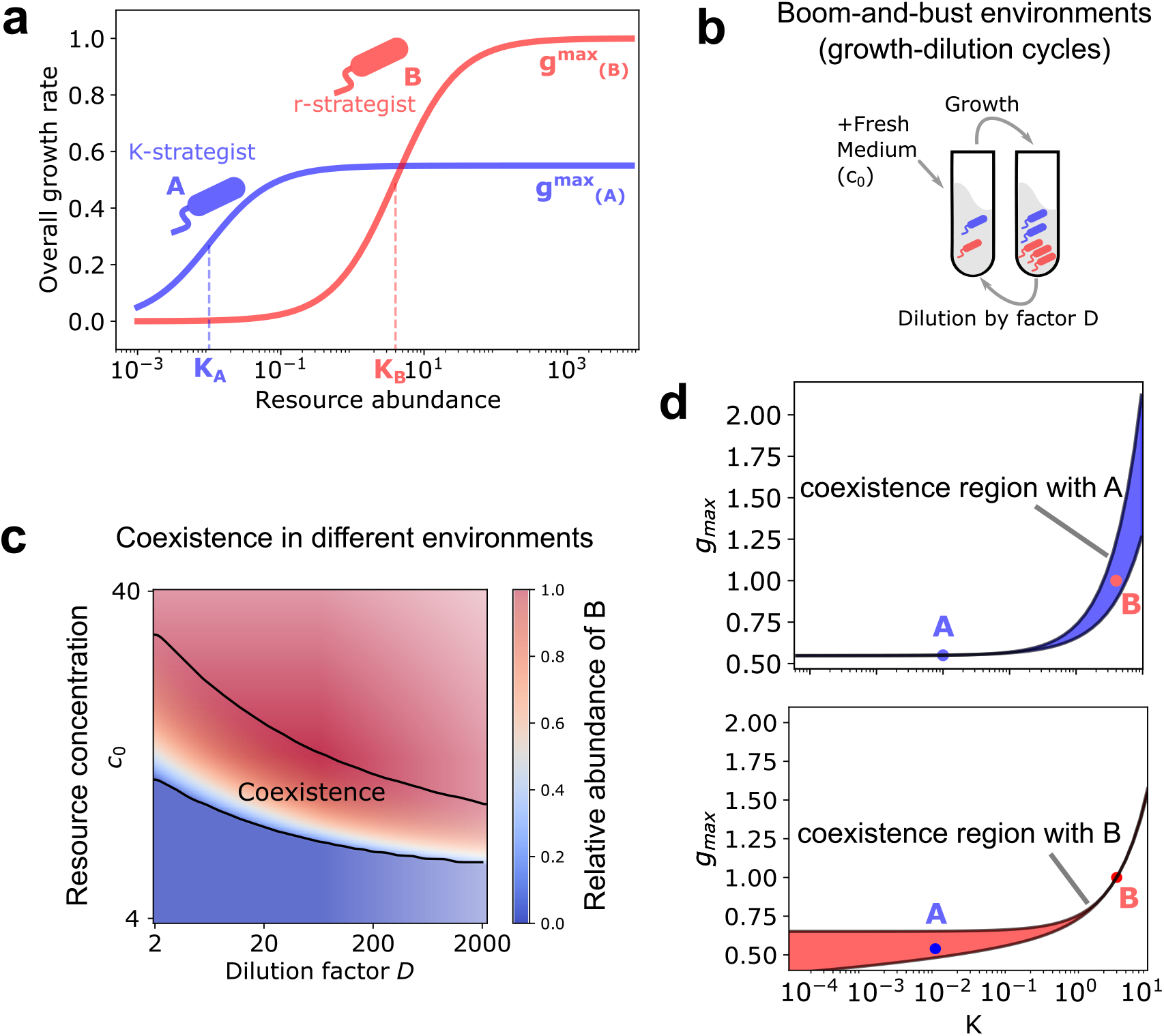
Coexistence of two strains in a boom-and-bust environment supplied with a single limiting resource. **(a)** Strains A and B follow Monod’s law of growth. Strain A is a *K*-strategist with a lower maximum growth rate *g_max_* and substrate affinity(*K*) compared with strain B, which is an *r*-strategist. **(b)** Schematic of the serial dilution experiment. At the end of each cycle, we dilute the population by a factor *D,* and supply fresh medium with resource concentration *c*_0_ (see Methods). **(c)**Feasible environmental parameters for two strains with given growth rates. Each point on the heatmap defines an environment, characterized by *c*_0_ and *D*, and the color of each cell represents the relative abundance of strain B at steady state. Black lines show the range of environments where both strains, A and B, coexist. **(d)** Coexistence region in terms of growth parameters of strain *B* for a fixed environment (D = 10, *c*_0_ = 10) and fixed strain *A.* The blue region represents the range of growth parameters of B (red dot) where it can coexist with strain A (blue dot) in a fixed environment. Black solid lines are theoretical boundaries of the region.

Simulations with two strains showed that both *r* and *K* strategists can coexist over a wide range of parameters *c*_0_ and *D* (Fig. 1c), as observed in previous studies (e.g., Supplementary Figure 13a in ref. [24] and Figure 6 in ref. [21]). To quantify the range of environmental parameters in which both strains could coexist, one can measure area of the space of parameters *c*_0_ and *D* where both strains survive (the region between the black lines in Fig. 1c).

Alternatively, one can fix the environmental parameters *c*_0_ and *D* and quantify the range of growth parameters of strain *B* which would allow it to coexist with strain *A* with a given set of growth parameters, *g_A_* and *K_A_*. This range, visualized in blue in Fig. 1d, is extremely narrow when *K_B_* is below or slightly larger than *K_A_*, but expands significantly as *K_B_* starts to increase compared with *K_A_*. Analytical calculations show that the width of the region, Δ*g*_coexist_ bounded from above by 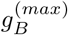 and from below by 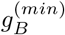, for which coexistence is possible, can approximately be written as follows (see Supplementary Text):

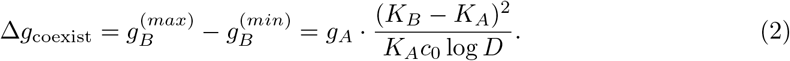

Here we assume that (*K_B_* – *K_A_*) ≪ *K_A_*. In a more general case, one should replace (*K_B_* – *K_A_*)^2^/*K_A_* with 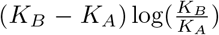 (Supplementary Text). Note that the width of this region depends only mildly (logarithmically) on the dilution factor *D* compared with its strong, inverse dependence on the resource concentration *c*_0_, suggesting that coexistence of strains should be possible for a broad range of environmental fluctuations.

In Fig. 1e, we show the complementary range of coexistence for a strain *B* (red in Fig. 1e), which lay inside the coexistence range of strain *A* (Fig. 1d). Note that coexistence range of *B* is broad for values of *g_A_* and *K_A_* below the growth parameters of *B*, and narrow on the opposite side (*g_A_* and *K_A_* above *g_B_* and *K_B_*, respectively). This is in contrast with the coexistence range of *A*, which is instead broader on the right.

Thus, coexistence between two strains *A* and *B* is possible in this environment because *B* can consume and grow on low concentrations of the available resource, on which strain *A* grows relatively poorly. In such boom-and-bust environments, “satellite niches” appear, enabling coexistence between *A* and other strains, which must lie in the region shown in Fig. 1d. In the rest of this manuscript, we will refer to this coexistence region for strain *A* as its “shadow”.

### Coexistence of three strains

Now we discuss the circumstances in which a third strain, *C*, can coexist with two already coexisting strains *A* and *B* (Fig. 2a) Note that strain *C* has an intermediate growth rate and affinity, i.e., *g_A_* < *g_C_* < *g_B_* and *K_A_* < *K_C_* < *K_B_*. Naively, one might assume that any whose growth parameters *g_C_* and *K_C_* lie within the shadow of the two other strains *A* and *B*, would be able to coexist with them. However, simulations of our model with three strains revealed that this is not the case (Fig. S1); instead the region where all three strains truly coexist with each other (shown in dark grey, Fig. 2b) is a smaller sub-region of the intersection of both pairwise shadows (Fig. S1). In hindsight, this is not surprising. Indeed, two strains lying in the shadow of each other only ensures their pairwise coexistence. Generally, pairwise coexistence between all pairs does not guarantee that all three strains will be able to simultaneously coexist.

**Figure 2:**
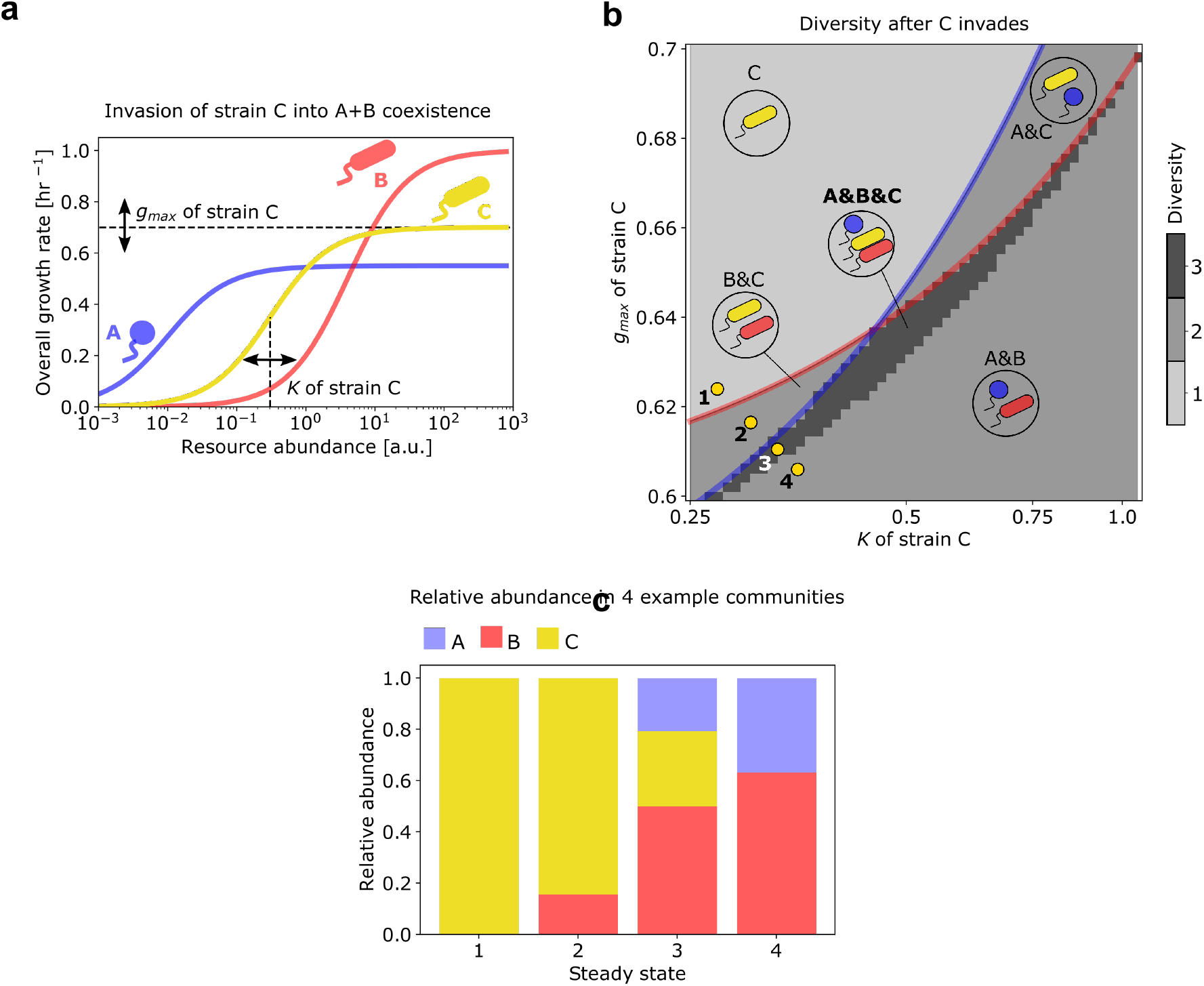
Three-strain coexistence in a boom-and-bust environment with single resource. **(a)** Growth rate profiles of strains *A, B* and *C*, which all obey Monod’s law of growth. We fixed the *g_max_* and *K* for *A* (blue) and *B* (red) fixed, while varying them for strain *C* in a manner that both parameters *g_C_* and *K_C_* remained between those of strains *A* and *B*. **(b)** Heatmap showing the final community diversity (number of strains) as a function of the growth parameters *g_C_* and *K_C_*. Given a strain C with variable *g_max_* and *K* along with fixed strains A and B, the heatmap shows the number of coexisting species in the steady state of serial dilutions. Here, the region above the blue line represents where C drives A out of the steady state, and the region above the red line represents where B is driven out by C. The yellow dots stand for 4 possible strain C’s that will result in 4 different steady state compositions. **(c)** Possible compositions of the final community. For 4 mutants of C represented by yellow dots in (b), bar plots show the relative abundance of the 3 strains in the steady state.

An important question to ask about strains occupying satellite niches is whether their abundances will be much lower than the strain with the highest maximal growth rate *g* (here, strain *A*), occupying the key resource niche. By tracking the abundance distributions of communities within and near the shadows of the three strains, we found that when multiple strains coexist, their abundances are not greatly skewed towards strain *A*, instead being comparable to each other (Fig. 2c). Specifically, as we sample communities with growth parameters *g_C_* and *K_C_* across a line shown in Fig. 2c, the community structure changes from being exclusively consisting solely of strain *C* (Fig. 2b, point 1), to coexistence between *B* and *C* (point 2), to all three strains coexisting (point 3), and finally to point 4, where strains *A* and *B* coexist, but *C* goes extinct.

In the case of strains following Monod growth, obtaining the boundaries of the region where three or more strains can coexist is mathematically challenging, because of which we resorted to numerical simulations with a large pool of strains. However, we also studied a simplified nonlinear growth law where the boundaries of the coexistence region for any number of strains could be computed exactly (see Supplementary Text, equation S35 for 3 species and S39 for an arbitrary number of species, *n*). Microbial growth in this simplified version of the model approximates Monod’s law by a step-like function, with the growth rate being zero below resource concentration *K*, and *g* above concentration *K*. In principle, an extension of this simplified model to include multiple steps can approximate any growth law to arbitrary precision, provided a sufficiently large number of steps. This extended model with multiple steps is mathematically equivalent to another model [25] where multiple microbial strains grow in a diauxic manner, starting from their most preferred resource, switching to progressively less preferred resources, being depleted one resource at a time. Each diauxic shift in this extended model would occur upon a small decrease in concentration of the primary resource *C*. Note that in this mathematical formulation, all microbial strains use resources in the same order of preference, thereby making coexistence harder to achieve as the number of strains increases. In fact, a previous study showed that coexistence in diauxic communities is most likely when resource preferences are complementary to each other, and least likely when they are identical [9, 25].

### Statistics of the number of coexisting strains

To investigate how many strains can typically coexist in such environments, we simulated community dynamics of a large randomly generated pool of 100 strains (Fig. 3a), whose growth parameters *g* and K were sampled from suitably chosen probability distributions (individual points in Fig. 3c; Methods). In over 2,000 such simulations initialized with different species pools, we carried out community assembly simulations (Methods). The most common outcome in these simulations was a community with just one strain (96.6% of cases), however in 3.35% of simulations, we did observe 2 strains and in the remaining 0.05%, 3 strains. The likelihood of observing *n* strains decreased approximately exponentially with n. In fact, our simulations could have yielded communities with even more than 3 strains, but given the expected rarity of that occurring, it would not be a pragmatic use of computational resources. The observation that with a reasonable species pool size and number of simulations, one obtains no more than 3 strains coexisting, is in agreement with results from Stewart and Levin [21]. The surviving strains (Fig. 3c, dark red points) in our three-strain communities had among the highest *g* and lowest *K* values, and appeared to lie on a Pareto frontier. However, predicting exactly which of the strains on the Pareto frontier will survive is challenging.

**Figure 3:**
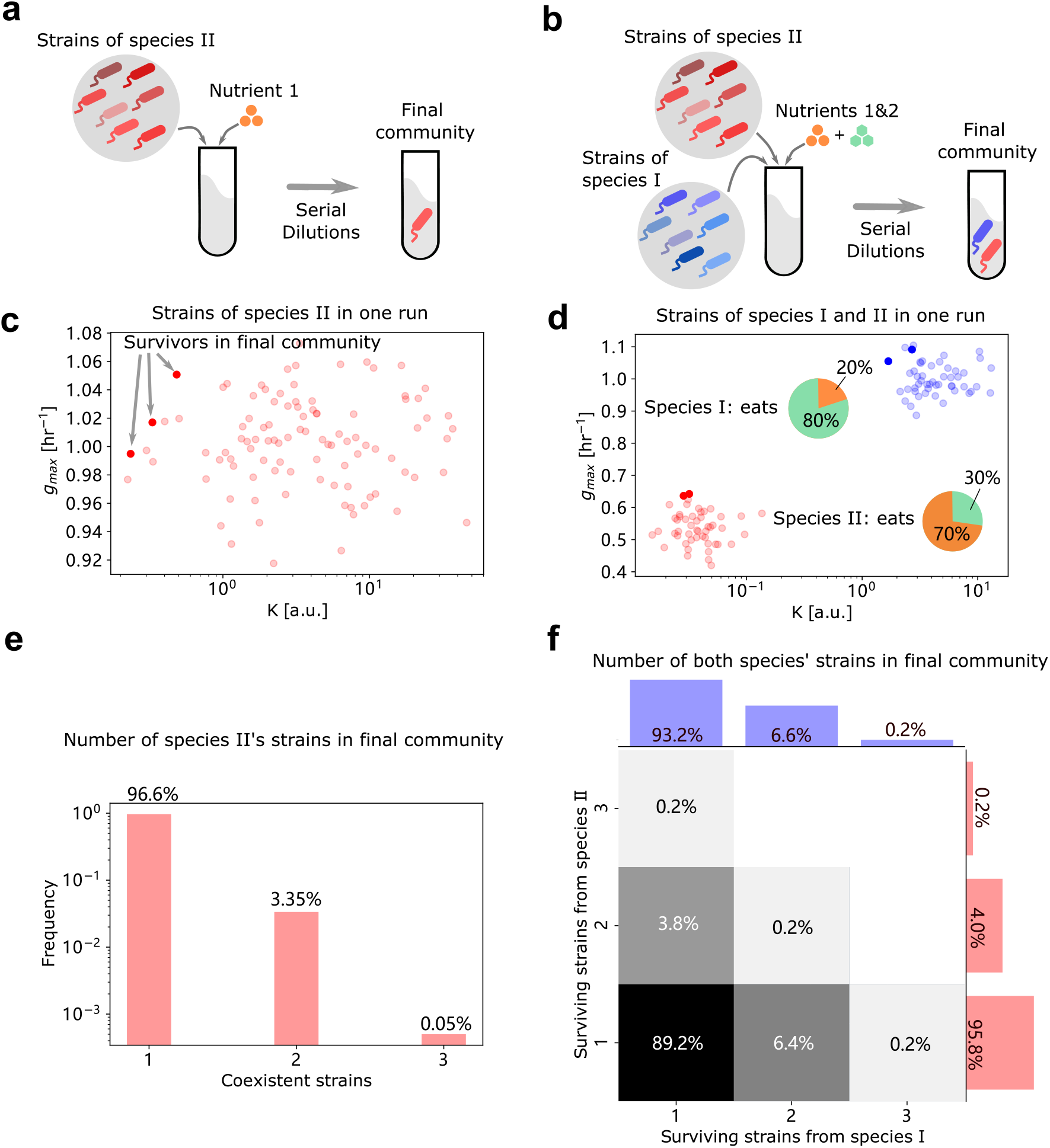
Final communities from strain pools. Panel (a)(c)(e) show results from single species in a single resource environment, and (b)(d)(f) depict results from 2 species in a 2-resource environment. **(a)-(b)**Schematic of strain pool simulations. All randomly generated strains from each species are presented together at the beginning of each serial dilution simulation, which will reach a final community at steady state. **(c)-(d)**Examples of strains in a strain pool. Dots show the *g_max_* and *K*’s of each strain, where the deep-colored ones represent the strains that survive in the final community. Blue dots are strains from species I and red from species II. Pie charts in (d) show the co-utilization profiles of each species. **(e)-(f)**Histograms of number of survivors in the final community. Bar plots of blue and red shows correspondingly the number of surviving strains from species I and II. Heatmap in (f) separately represents the distribution of 2 species’ number of survivors.

### Extending results to complex communities and multi resource environments

Until now, we showed that multiple strains specializing on only one limiting resource could coexist with each other by using different *r* or *K* strategies. In natural communities, however, strains will not only have access to multiple resources, but also face competition from multiple other species. Indeed, different species might have evolved different resource preferences, lowering niche overlap and increasing the probability of coexistence with each other. In light of this, we next investigated whether our proposed mechanism of strain coexistence — each with different *g* and *K* parameters — was robust to the presence of multiple resources and species. As the simplest such extension of our previous results, we simulated community assembly of strains from two distinct species, I and II, on two different substitutable resources, *R*_1_ and *R*_2_. Each species co-utilized both resources, but each had a different preference towards one of the two resources: namely, species I preferred *R*_1_, while species II preferred *R*_2_. We also assumed that all strains of the same species had identical resource preferences, but different growth parameters, g and K, on each of the two resources (see Methods). We then simulated community assembly via serial dilutions, each starting with a different pool of randomly generated strains belonging to two species, whose resource preferences were fixed (Fig. 3c–d). At the end of several growth-dilution cycles, we asked how many strains of each species could coexist in the final, steady state communities.

Fig. 3d shows the outcome of one such community assembly simulation, with 50 strains belonging to each species (shown as red and blue points in Fig. 3d). Species I, which preferred resource *R*_1_, dedicated 80% of its consumption (quantified by the growth rate coefficient *g/K* at low resource concentration) to *R*_1_ and 20% to *R*_1_; species II instead dedicated 30% of its consumption to *R*_1_ and 70% to *R*_2_. At steady state, two strains of species I (dark blue points in Fig. 3d) and two strains of species II (dark red points in Fig. 3d) coexist in the community. The growth parameters *g* and *K* of these strains are positioned at the Pareto frontiers (upper-left corners of the *g* – *K* plane) of each of their species pools.

By performing community assembly simulations over 500 independently generated species pools, we observed that in 89.2% of cases, both species could coexist, but only with one strain in each species (Fig. 3f). In the remaining 10.8% of cases, at least one species was represented by > 1 strain. The blue and red bars in Fig. 3f show the marginal distributions of the number of strains observed in each of the two species (Fig. 3f). Note that each of the marginal distributions bears striking resemblance to the histogram in Fig. 3e, where we simulated only one species and one resource. Thus, our results about the coexistence of multiple strains of the same species are robust to the inclusion of more than one resource (or niche) occupied by different species.

## Discussion

In this paper, we showed that in time-varying environments, the competitive exclusion principle can be broken through the formation of a few satellite niches alongside the primary resource niche. These niches can be occupied by closely related strains belonging to the same species, which rapidly diversify in two key parameters, i.e., their maximum growth rate and resource affinity. Our results add to a substantial body of work investigating how nonlinearities in resource-dependent growth can lead to violations of the competitive exclusion principle [26, 21, 27]. As in these studies, in our work satellite niches only appear in time-varying environments, such as the boom and bust cycles we investigated here, and completely disappear in static environments, such as chemostats. In chemostat-like static environments, resource concentrations and microbial abundances reach a fixed value at steady state, resulting in strict competitive exclusion, where only 1 strain can survive per resource (primary niche). The main property of time-varying environments that create satellite niches is that resource concentrations change with time, creating opportunities for different strains to specialize and be more competitive in different concentration ranges. In environments with more than one supplied resource (primary niches), species could also specialize to change the ratio in which they consume different resources, i.e., change how they allocate their enzyme budgets. Following in the footsteps of Good et al. [28], we adapted our model to include the possibility of small variations in resource budget allocation by closely related strains. We found that variation in budget allocation alone rarely (1% of simulations) leads to coexistence of multiple strains, compared with variation in growth parameters alone (11.8% of simulations, Fig. S2). Thus, satellite niches created by boom and bust cycles chiefly select for variation in such growth parameters.

The existence of organisms which specialize either on rapid growth (*r* strategists) or more complete depletion of the available resources (*K*-strategists) is well-established in natural ecosystems [22, 23, 29]. Examples include microbes residing in Earth’s upper ocean, where *r*-strategists dominate in strongly seasonal temperate oceans, while *K*-strategists dominate in stable, low-seasonality, equatorial marine environments [30, 31]. However, whether such *r* and *K* strategists can coexist in the same environment has not been fully explored. Our results show that *r*- and *K*-strategists may indeed coexist as closely related strains of the same species, both in simple and complex multi-species, multi-resource environments. These predictions also match spatial distributions of microbial populations in Earth’s upper oceans predicted in ref. [32]. While the strain diversity in highly seasonal environments (up to 8) was somewhat higher than predicted by our model (up to 3), the authors readily admit that their predictions might be somewhat inflated by ocean dynamics, transiently mixing organisms from different habitats. Furthermore, rapid and continuous evolution might also increase the apparent diversity of a community, due to transient strains generated by mutations and ultimately lost due to competitive exclusion. Our model included a fixed pool of strains and provided sufficient time to achieve a steady state as a result of competitive exclusion, and thus ignored the transient diversity due to rapid evolution.

Another natural environment in which limited strain diversity has been observed is the human gut microbiome [7], where the coexistence of a few strains (between 1 and 3, termed “oligo-colonization”) was widely observed. In particular, the authors mention that “it is not clear what mechanisms would allow for a second or third strain to reach intermediate frequency, while preventing a large number of other lineages from entering and growing to detectable levels at the same time”. Our work, in quantitative agreement with these observations, suggests that a possible mechanism explaining them might be that closely related strains differ in their growth parameters (*g_max_* and *K*) by which they consume resources.

“Oligo-colonization” is not limited to natural environments such as ocean and gut microbiomes, but also manifests in microbial communities domesticated in the lab [15]. In these domesticated communities, between 1 and 2 closely related strains of the same species were found to coexist over multiple (~ 70) boom-and-bust (serial dilution) cycles, consistent with the predictions of our model. Thus, taken together, our model explains a possible mechanism by which such “oligo-colonization” of a small number of closely related strains might be widespread in both natural and laboratory-domesticated microbial communities.

## Acknowledgements

We thank Maria Verbitsky for writing the Python code to find the environmental conditions in which any two strains may coexist. We thank the Kavli Institute for Theoretical Physics, UCSB, where this project took root, and Daniel Fisher for discussions which helped lead us to the analyses in this paper. This research was supported in part by the National Science Foundation under Grant No. NSF PHY-1748958, as well as NSF Grant No. PHY-1748958 and the Gordon and Betty Moore Foundation Grant No. 2919.02. A.G. is supported by the Gordon and Betty Moore Foundation as a Physics of Living Systems Fellow through Grant No. GBMF4513.

## Methods

### Serial dilutions

In our serial dilution simulations, we present every strain at equal amount of 1 unit at the beginning. In each dilution cycle, *c*_0_ = 10 units of every resource is added into the system, and strains grow according to Monod’s law for *T* = 24 hr. At the end of each cycle the system is diluted by a factor of *D* = 10.

### Strain coexistence

We use two example strains (A and B) to investigate their coexistence in a single resource environment. Their maximum growth rates are 0.55 hr^-1^ and 1.0 hr^-1^ respectively, with substrate affinity being 0.01 a.u. and 4.0 a.u. In Fig.2b, we introduced an invader strain C whose maximum growth rates vary between [0.6, 0.7] hr^-1^ and substrate affinity vary between [0.25, 1.05] a.u.

### Strain pool simulations

For the single resource simulations, we generated 2000 strain pools of species II independently, each containing 100 mutant strains. Their maximum growth rates were sampled independently from a normal distribution of 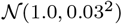 in hr^-1^. Their substrate affinities were sample independently from a log-normal distribution of logNormal(log 4, 0.5^2^) in a.u.

For the 2-resource simulations, we generated 500 strain pools independently, each containing 50 mutant strains from species I and 50 from species II. Both species co-utilize the 2 resources, and as an example, the overall growth rate of strain A is

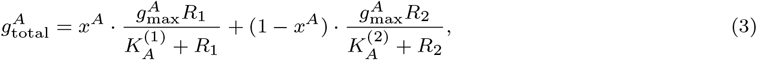

where *x^A^* and 1 – *x^A^* are coefficients of co-utilization.

The coefficient of *x* is fixed at 0.2 for species I and 0.7 for species II. The maximum growth rates of mutant strains of species I and II are sampled from normal distributions 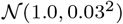 and 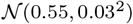 respectively. Strains of species I have their substrate affinities sampled from log-normal distributions: 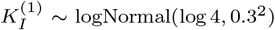, 0.3^2^), and 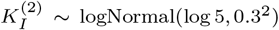. Strains of species II also have their substrate affinities sampled from log-normal distributions: 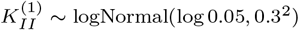, and 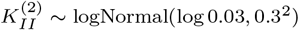.

In Fig. S2, we also investigated the case where co-utilization coefficients *x* is variable. They are independently sampled from normal distribution, 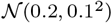 for species I and 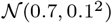 for species II.

## Code availability

All code comprised of custom Python scripts, using Python v3+, along with the Numpy v1.20.1 and Scipy v1.6.2 packages, and can be found at https://github.com/maslov-group/Coexistence_of_g_and_K.

## Data availability

All the numerical data from simulations can be found on the GitHub repository, at the following link: https://github.com/maslov-group/Coexistence_of_g_and_K. Source data are provided with this paper. No new experimental data was generated during this study.

## Supplementary Text

### 1 Monod’s growth allows for the satellite niche formation

#### 1.1 Serial dilution vs constant nutrient supply: why, in the former case, the competitive exclusion principle can be violated

The competitive exclusion principle is usually thought to restrict the number of species surviving in the environment to be less or equal to that of the unique food sources. (Barring crossfeeding and other types of interaction which can be understood as providing additional, positive or negative, sources.) However, it is also known that the species growing according to the Monod’s law can be either “*r*-strategists” maximising their growth rate on a given source when it is supplied in plentiful amounts or “*K*-strategists” striving to survive on less amounts of food whenever the supply is scarce. While in the case of the constant nutrient supply (chemostat) the number of species that grow under the Monod’s dynamics can never exceed the number of (generalized) sources, the environments that resemble the serial dilution experimental setting under certain supply regimes may allow for the survival of more than one species on a single resource.

This can happen because the Monod’s dynamics, under which the logarithmic growth rate for a species *B_i_* is given by *g_i_ c*/(*c* + *K_i_*) (with *g_i_*, *K_i_*, *c* being the Monod’s parameters of the microbe’s and the nutrient’s current abundance, respectively) may favor, say, first *B*_1_, when the nutrient is still comparatively abundant, and then *B*_2_ when the nutrient is closer to being depleted. When we are talking serial dilution, it is important for a species’ successful invasion to make sure that its abundance gets multiplied at least *D*–fold during a single dilution cycle (*D* being the dilution coefficient). Comparing the two logarithmic multiplication rates we see that, if *g*_1_ > *g*_2_,

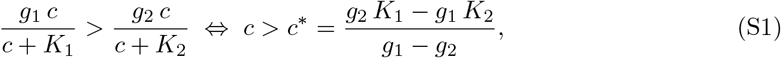

so that at *K*_1_/*K*_2_ > *g*_1_/*g*_2_ there is a turning point *c* = *c*\*. When, during the dilution cycle, the nutrient concentrations drops below that point, *B*_2_ starts multiplying faster than *B*_1_ and thus, in total, under some conditions may make it *D*–fold simultaneously with *B*_1_. No degeneration in terms of g, *K*–parameters is required for the coexistence to be achieved: even if narrow, it is still not a null-dimensional curve in the *g*, *K*-plane, but rather a region of parameters.

#### 1.2 Coexistence of two species on a single source: the criterion

Now we proceed at deriving the formal criterion of the necessary and sufficient conditions under which two microbes obeying the Monod’s growth law can coexist on a single source in a serial dilution setting.

Consider a serial dilution setting. Let the resource *c* be supplied in the amount *c*_0_ in the beginning of each dilution cycle, long enough for it to get effectively depleted in the end by the microbe or microbes invading the environment. Let the bacteria *B*_1_ with the maximum growth rate *g*_1_ and the *K*–parameter *K*_1_ establish a monoculture steady state in this environment. During each dilution cycle, the microbe will grow, so that

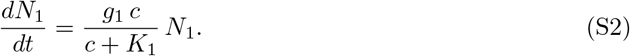

The resource c gets depleted as the microbe grows. For the sake of simplicity we set the yield equal to 1, so that

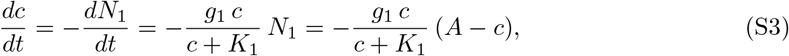

where the constant *A* comes from the conservation law (valid, separately, for each dilution cycle):

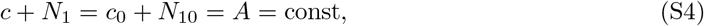

*N*_10_ being the initial abundance of the microbe at the beginning of a dilution cycle.

**Figure S1:**
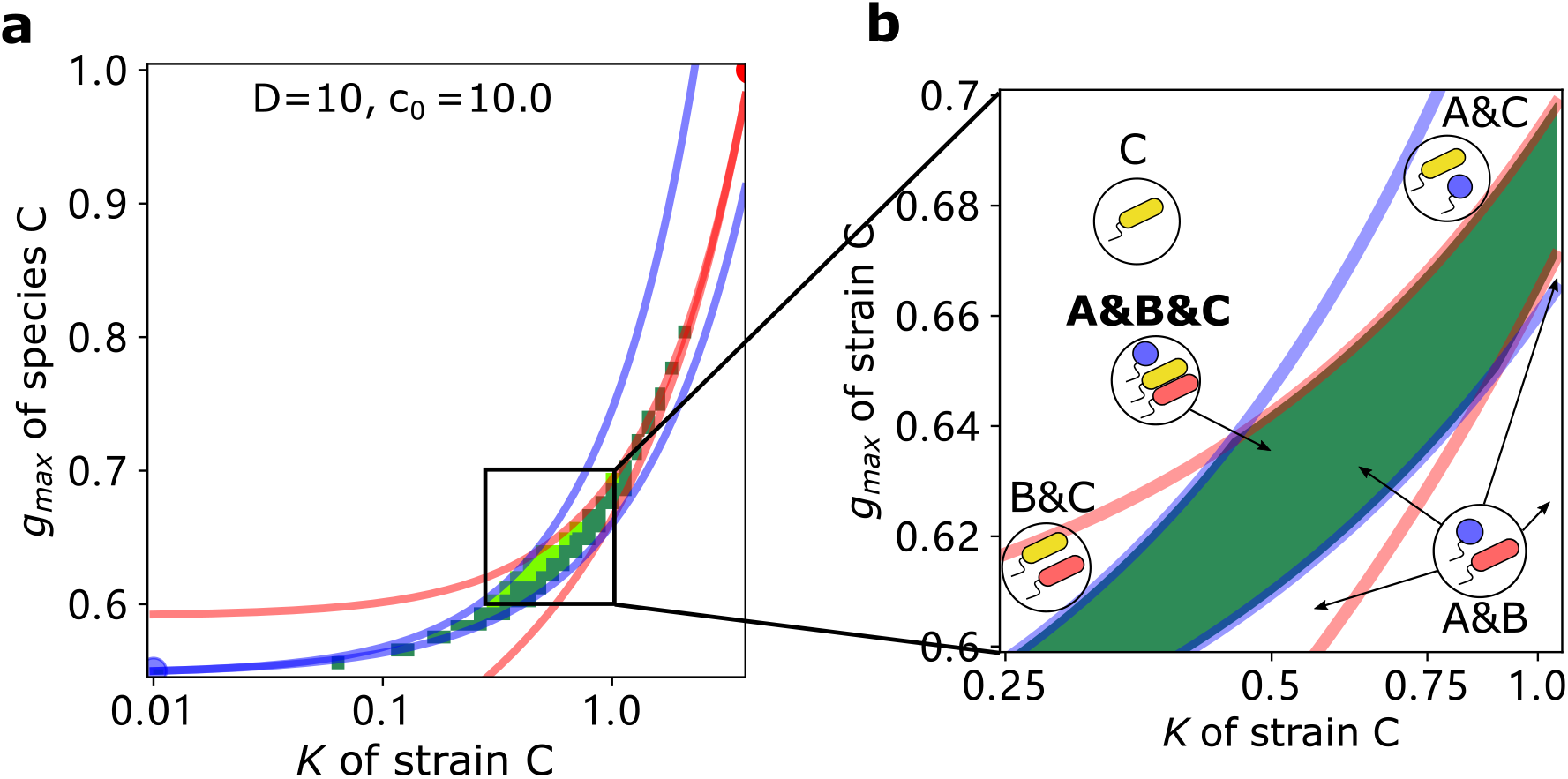
A detailed view of multi-strain coexistence, similar to Fig. 2b.

**Figure S2:**
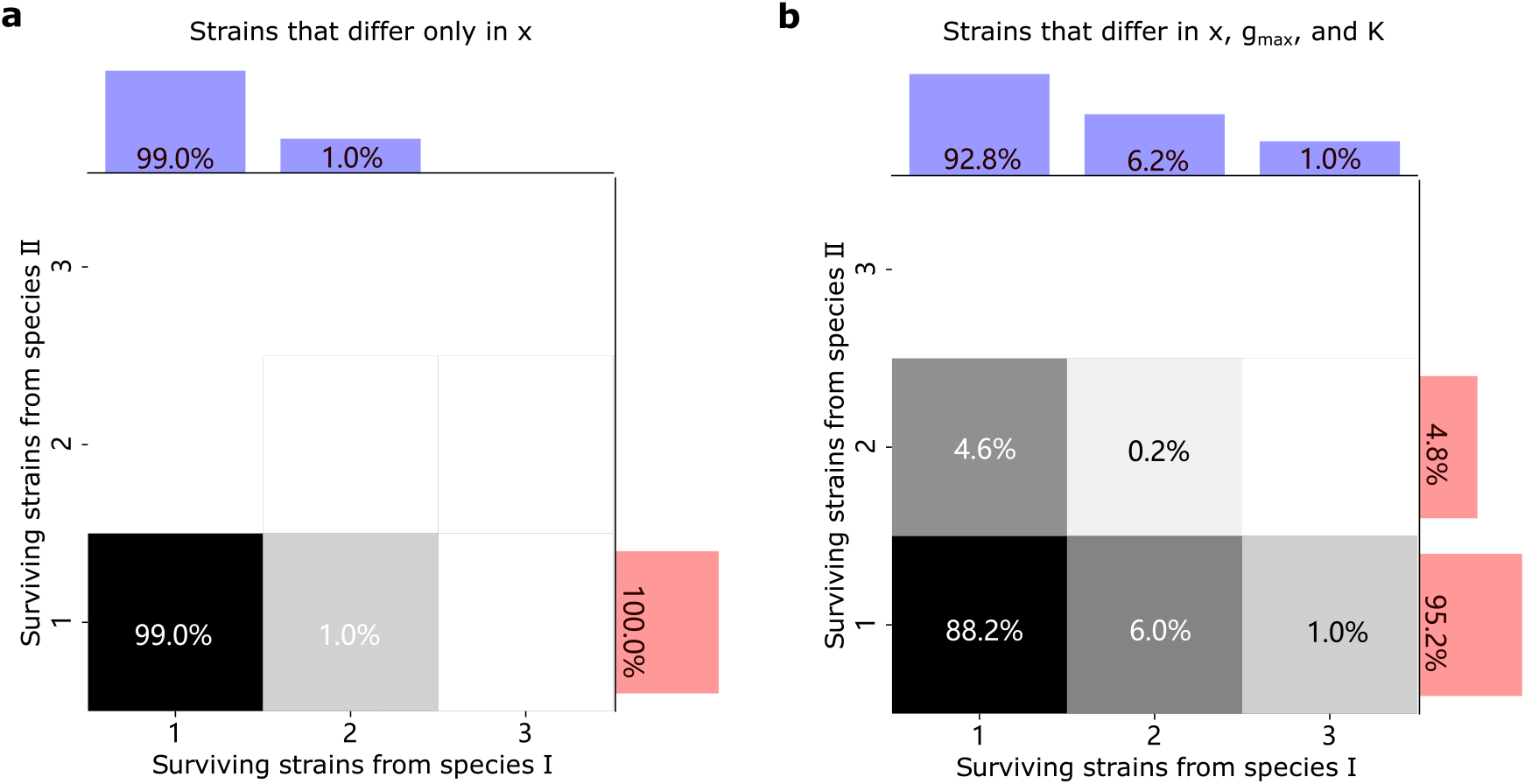
Histograms showing statistics of the number of coexisting strains: (a) when we allow small variations in how a species partitions two resources, with no variation in Monod growth parameters, and (b) when we allow changes in both the Monod parameters as well as resource partitioning. Bar plots on the margins in blue and red show the number of surviving strains from species I and II respectively. In (a), the variation in resource partitioning was as follows: each species’ resource partitioning was independently sampled from a normal distribution, 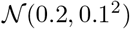 for species I and 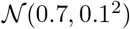 for species II.

If the dilution cycle is long enough, then for the steady monoculture state we can write effectively

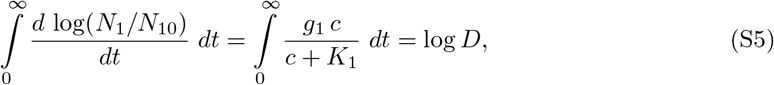

as the multiplication coefficient (the number by which the initial abundance is multiplied towards the end of each dilution cycle) should be equal to the dilution coefficient *D*. For the same reason the steady state initial abundance of the microbe for each dilution cycle should be

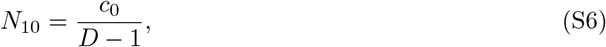

as can be easily seen using (S4). Thus the constant *A* appearing in (S4) in the steady state is found to be equal

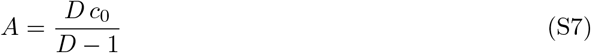

If a microbe *B*_2_ is able to invade the monoculture environment (shaped by *B*_1_) in the seed amount, then through its first dilution cycle in the environment it should grow more than *D*–fold.

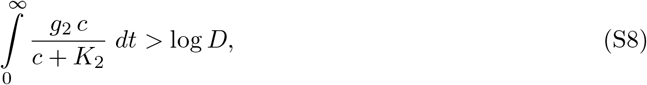

As, upon introduction into the environment, the invader can influence the dependence *c*(*t*) in but a negligibly small way, we can use (**??**) to give *dc*/*dt* as we rewrite (S8) changing variables and going from integration over *t* to integration over *c*:

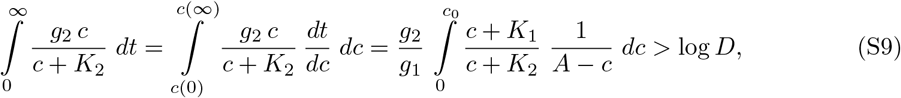

which, upon integration, together with (S7), yields the condition under which the species *B*_2_ can invade the monoculture environment shaped by *B*_1_:

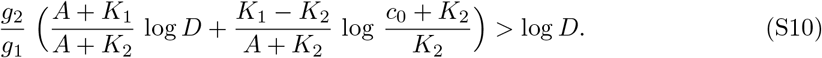

The condition under which *B*_1_, in its turn, can invade the monoculture environment shaped by the microbe *B*_2_ is obviously obtained from (S10) by swapping the indices 1 and 2. Thus, for a given *B*_1_ with the Monod parameters (*g*_1_, *K*_1_) for each *K*_2_ < *K*_1_ there is a range from which *g*_2_ can be chosen so that *B*_1_ and *B*_2_ can share the environment. (Indeed, if both *B*_2_ can invade the monoculture of *B*_1_ and *B*_1_ can invade the monoculture of *B*_2_, no one of the microbes will be able to outcompete the other, so that they would have to share and coexist.) The said range is given by the following:

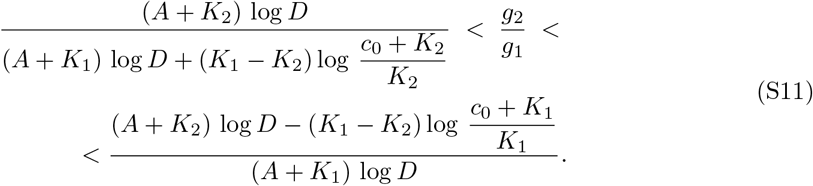

#### 1.3 The “mutation-friendly” limit: two strains of the same species (close to each other in terms of their growth rates) coexistence criterion

In the limit *c*_0_ ≫ *K*_1_ > *K*_2_ and reasonably large values of *D*, say, *D* ~ *c*_0_/*K*_1_ the microbes sharing the same niche may do so only if they are very close in terms of their growth rates, and those are neatly tuned to each other. If there is a biological reason for a species to be able to lessen considerably its *K*–value at the expence of decreasing its own growth rate by a tiny little bit, one can observe two strains of the same species surviving on a single source in an environment with boom and bust cycles. Indeed, in this limit *A* = *Dc*_0_/(*D* – 1) ≃ *c*_0_, *c*_0_ + *K*_i_ ≃ *c*_0_ for *i* = 1, 2, so that the coexistence criterion (S11) can be reduced to

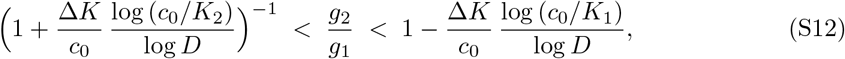

Where Δ*K* = *K*_1_ – *K*_2_. As, on the left side of the equation above, the second term is small compared to 1, one could proceed with the reduction to get

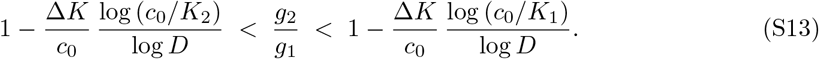

Thus, in this limit, the coexistence range for the value of the ratio *g*_1_/*g*_2_ is thin, its width *r* given by

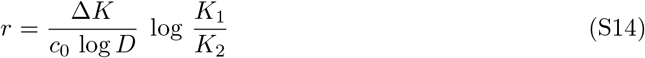

The larger is *K*_1_/*K*_2_, the wider the range. On the other hand, if *K*_1_ – *K*_2_ = Δ*K* → 0, one gets log(*K*_1_/*K*_2_) = log(1 + Δ*K*/*K*_2_) ≃ Δ*K*/*K*_2_, so that for the strains very close to each other in both *g* in *K* we get:

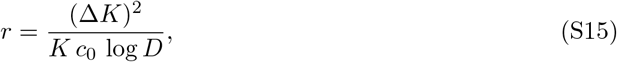

with *K* ≃ *K*_2_ ≃ *K*_1_. (It can be shown that the same quadratic dependence of the *g*_2_/*g*_1_ range width on Δ*K* at Δ*K* → 0 holds true for any values *D* > 1 and *c*_0_.)

For a given microbe *B*\* with the Monod parameters (*g*\*, *K*\*) the species that can coexist with it for any given initial resource concentration *c*_0_ form a banana-shaped shadow in the *g*, *K* plane (see Fig. 1d). So in an environment resembling a serial dilution setting rather than a chemostat, whenever the nutrient supply come somewhere in between “feast” and “famine”, each niche occupied by a specialist would be accompanied by a shadow of satellite niches, that can be occupied by yet another microbe, or, in principle, even by several other species, as we have shown in this paper.

#### 1.4 Why the coexistence range always exists

It remains to prove that the coexistence range (S11) exists for any reasonable initial abundance *c*_0_ of the resource (we assume *c*_0_ > *K*_2_) and for any value of the dilution coefficient *D* > 1. More precisely, one can choose any value of the ratio for a pair of *K*–parameters of the Monod’s dynamics, *K*_1_ > *K*_2_, *x* = *K*_1_/*K*_2_ > 1, and any value of the initial abundance *c*_0_ = *y K*_1_ = *xy K*_2_ in the same *K*–units, and be certain to find the ratio *g*_2_/*g*_1_ of the maximum growth rates of the two species such that, with a given dilution coefficient *D* > 1, they would be able to coexist on that single resource *c* in the serially diluted environment. This ratio *g*_2_/*g*_1_ would be bounded from above and from below, and any value within the range given by (S11) will work.

If we prove that, we can be justified in expecting the satellite niches to eventually emerge under the favourable conditions in nature. In the laboratory, however, more practical question would be whether, for given bacteria *B*_1_ and *B*_2_, with the Monod parameters (*g*_1_, *K*_1_) and (*g*_2_, *K*_2_) respectively, the appropriate conditions (i. e. the initial resource concentration *c*_0_ and the dilution coefficient *D*) can be found, so that they may share the environment. The answer seems to be positive if *g*_1_/*g*_2_ < *K*_1_/*K*_2_, as we will show later. Now we proceed to proving that a satellite niche can always emerge.

It is worth noting that, with *γ* = *g*_2_/*g*_1_,

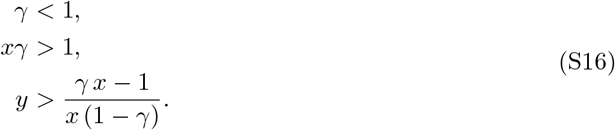

where the first inequality comes from our assumption that *g*_2_ < *g*_1_, while the second and the third follow from Eq.(S1) and the fact that *c*_0_ should be greater than the critical value *c*\* mentioned in (S1) so that the competing microbes could take turns in growing faster than their counterpart.

The said range will exist if, and only if,

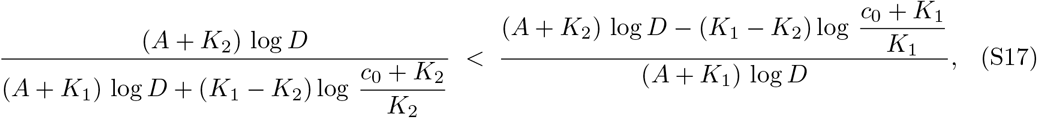

meaning simply that the lefthand side of the criterion inequality (S11) is indeed less than its righthand side.

Using *x, y* introduced above (so that the biomass conservation constant *A* = *Dc*_0_/(*D* – 1) = (*D*/(*D* – 1)) *x y K*_2_), we transform the inequality (S17) into

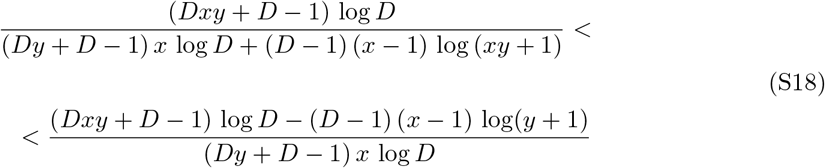

This is easily transformed by algebraic means into an inequality:

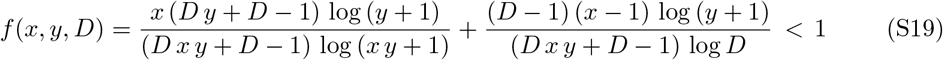

Within the relevant region of the parameters, it is possible to determine the maximum value of *f* (*x, y, D*) using elementary calculus. Namely, if we prove that *f*, at *x* > 1 decreases with *x* for any fixed values of *y* and *D* within the relevant region of parameters, and take into account that *f* (1, *y, D*) = 1 for any acceptable values of *y*, *D* which is easily checked by substituting *x* = 1 into (S19), then we prove the inequality in question.

Calculate

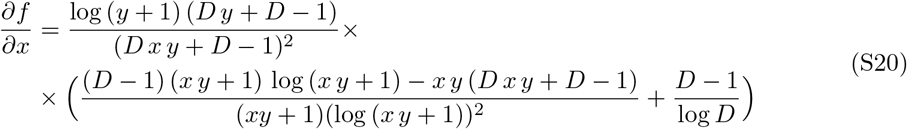

To determine the sign of *∂f*/*∂x*, one would have to concern oneself with the sign of the expression in the brackets, which we rewrite as:

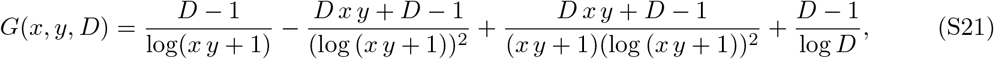

the only one of the factors constituting *∂f*/*∂x* that has a chance to change its sign (all the others are positive under the conditions imposed by the situation under consideration). Introduce *z* = *xy* + 1 (note that, by the virtue of (S16), *z* > 2) and rewrite the expression (S21) in terms of *z*:

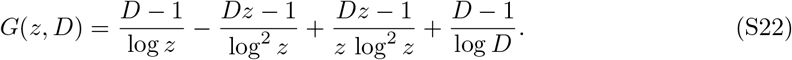

Differentiating (S22) in terms of *z*, we obtain

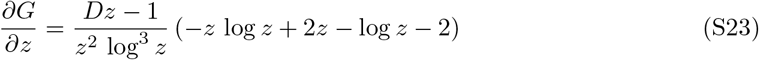

Now one should concern oneself with the sign of the expression

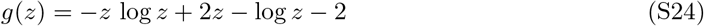

Differentiate it twice in terms of *z,* to find

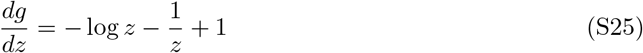

and

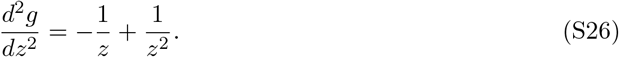

Now *d*^2^*g*/*dz* turns zero when *z* = 1 and is positive at *z* < 1 and negative at *z* > 1, thus *dg/dz* has its maximum at *z* = 1. Calculating

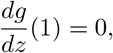

we see that at *z* > 1 the function *g*(*z*) decreases and thus must be always less than *g*(1) = 0, so that *G*(*z*), in its turn, decreases at *z* > 1.

Assume *z* = 1 + *ζ*, *ζ* > 0, *ζ* ≪ 1 and substitute in (S22). Upon expanding the Taylor series of log(1 + *ζ*) up to the required order of smallness, we get

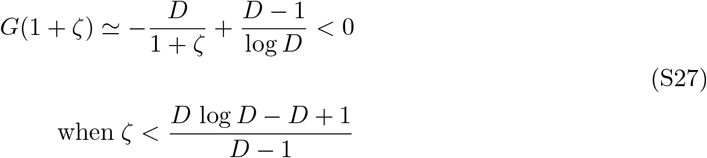

Thus at *z* → 1 we have *G*(*z*) < 0 and, since *G*(*z*) decreases with *z* when *z* > 1, we conclude that *G*(*z*) < 0 at *z* > 1. That is what makes *∂f* (*x,y, D*)/*partialx* from (S20) negative in the area we are interested in, and thus *f* (*x, y, D*) from (S19) at any fixed values *y, D* decreases at *x* > 1. Hence for any acceptable values of (*x, y, D*) we have *f*(*x, y, D*) < *f* (1, *y, D*) = 1 which proves (S19), and that, in its turn, is equivalent to having proved (S17).

Using elementary analysis we have proven that the inequality (S17) always holds and thus the range of *g*_2_/*g*_1_ fit for the coexistence always exists at any values of *c*_0_/*K*_1_ and *K*_1_/*K*_2_ > 1.

#### 1.5 What are the coexistence conditions for two given species?

As we have already noted, there is a question more relevant in a laboratory: given two species and a resource, what should the initial concentration *c*_0_ of this resource amount to, and which value of the dilution coefficient *D* should one take in order to get the bacteria steadily coexist in the serial dilution experiment?

Rewrite (S11) in terms of *x*, *y*, *γ*. As, again, *K*_1_ = *x K*_2_, *c*_0_ = *y K*_1_ = *x y K*_2_ and *A* = *D c*_0_/(*D* – 1) = *D x y*/(*D* – 1), we get

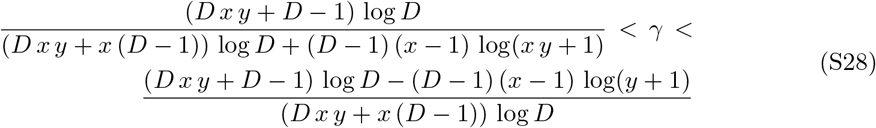

It is useful to remember that to enable coexistence one must have *c*_0_ > *c*\* > 0, *c*\* = (*g*_2_ *K*_1_ – *g*_1_ *K*_2_)/(*g*_1_ – *g*_2_) being the critical value of the initial concentration at which the “*K*–strategist” catches up with the “*r*–strategist” in terms of their growth rates. (Upon further depletion of the resource the “*K*–strategist” becomes the one that grows faster.) In terms of *x,y,γ* it’s

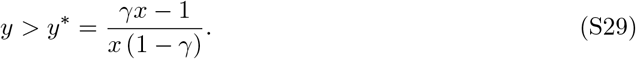

First it should be noted that at *D* → 1 the leftmost side of the inequality (S28) tends to 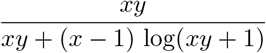. At *y* → ∞ this amounts to 1. To see why, consider *α*(*t*) = (log(*t* + 1))/*t* at *t* → ∞. Using the L’Hospital’s rule at *t* → ∞ we get *α* → 0. Dividing both the nominator and the denominator of l. h. s. of (S28) by *xy* and making *y* tend to ∞ we get 1/(1 + (*x* – 1) *α*(*xy*)), which tends to 1 as *α*(*xy*) → 0.

Thus at *D* → 1 and *y* → ∞ we can always make l. h. s. of (S28) greater than *γ*, which is less than 1, say, at some *D* = 1 + *ϵ* (with *ϵ* > 0) and *y* = *Y*.

On the other hand, at *y* → *y*\* = (*γx* – 1)/(*x* (1 – *γ*) and *D* = *xy* + 1 the l. h. s. of (S28) becomes

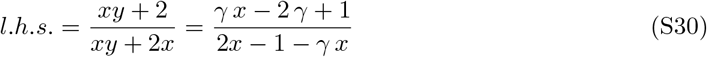

Comparing this with *γ* is easily reduced to comparing 0 with (*γ x* – 1)(1 — *γ*) > 0, so at *D* = *xy* + 1 and *y* → *y*\* (and thus at some *y* = *y*\* + *δ*, *δ* > 0) we get l. h. s. of (S28) less than *γ*.

A path in (*D, y*)–plane joining the points (1 + *ϵ*; *Y*) and *x* (*y*\* + *δ*) + 1; *y*\* + *δ*) by the virtue of continuity will have a point (*D*_0_; *y*_0_) at which l. h. s. = γ. As the r. h. s. of the same inequality (S28) is always greater than l. h. s., at that same point (*D*_0_; *y*_0_) r. h. s. > *γ*. Somewhere in the close vicinity of (*D*_0_; *y*_0_), again due to continuity, there exists a point (*D*_1_; *y*_1_) at which l. h. s. < *γ* and still r. h. s. > *γ*, so that *γ* can be indeed coaxed into the interval between l. h. s. and r. h. s. in some area of the (*D*, *y*–)plane. This, however, is true only provided that the basic requirements *γ* < 1, *xγ* = (*K*_1_*g*_2_)/(*K*_2_*g*_1_) > 1 are satisfied.

To provide the conditions (*c*_0_, *D*) favoring the coexistence of the bacteria *B*_1_ and *B*_2_ with the Monod parameters (*g*_1_, *K*_1_) and (*g*_2_, *K*_2_) respectively, we offer a code that inputs the Monod parameters and returns a plot picturing the area of (*c*_0_, *D*) favouring coexistence (or the error message if the basic conditions mentioned above are not satisfied). All the points in the plot are clickable, so that the desirable numerical values can be obtained from it.

### 2 Simplified model of Monod-like coexistence of multiple species on a single resource

The idea that the competitive exclusion principle might be violated in the serial dilution setting, once adopted, might play with the elementary algebraic associations one routinely exploits and thus has at hand. After all, the Monod’s dynamics is characterised by two parameters (*g, K*); it might seem natural that it should, in principle, produce two solutions corresponding to two survivors supported by a single shared resource. In fact, this kind of intuition here goes amiss. In principle, we could have any number of survivors on a single resource. (The range of parameters favoring such coexistence grows smaller with each new prospective survivor added, so that, in reality, it is almost impossible to arrange the coexistence of more than two or three species.)

To see why that happens, consider a simple Monod-inspired model. Let *B_i_* be a microbe species that grows exponentially on a resource c with the maximum growth rate *g_i_* as long as the resource concentration *c* > *K_i_*. Then it stops growing. However discontinuous, that would be still a dynamics that is characterised by two parameters *g_i_*,*K_i_*. Fig. (2) shows three species and their logarithmic growth rates as dependent on the resource concentration c.

We consider the same serial dilution setting as before, with the dilution coefficient c and the length of the dilution cycle long enough for the bacteria to consume as much of the resource as they can.

Just as in the case with the real Monod’s dynamics, it is easy to see that two microbes *B*_1_, *B*_2_ can coexist using the same single resource *c*, as long as, if *K*_1_ > *K*_2_, then also *g*_1_ > *g*_2_ (otherwise one microbe at any point will grow faster than the other, until both stop growing, and thus will become the sole winner). Indeed, in the steady state, when the point *R*_1_ = *K*_1_ is reached, the concentration *N*_1_ of the microbe *B*_1_ should grow *D*—fold compared to the initial concentration *N*_10_ at the beginning of a dilution cycle within the possibilities granted by the exponential growth, *N*_1_ = *DN*_10_. Thus this should happen at the time point *T*_1_ = (log *D*)/*g*_1_. By that time the microbe *B*_2_ growing exponentially with the rate *g*_2_ will have the concentration *N*_2_ = *D*^*g*_2_/*g*_1_^ *N*_20_, where *N*_20_ is the initial concentration of the microbe *B*_2_ at the beginning of a dilution cycle in the steady state. The biomass conservation law at this point will read as

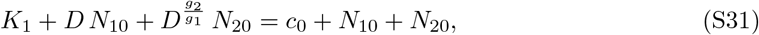

**Figure S3:**
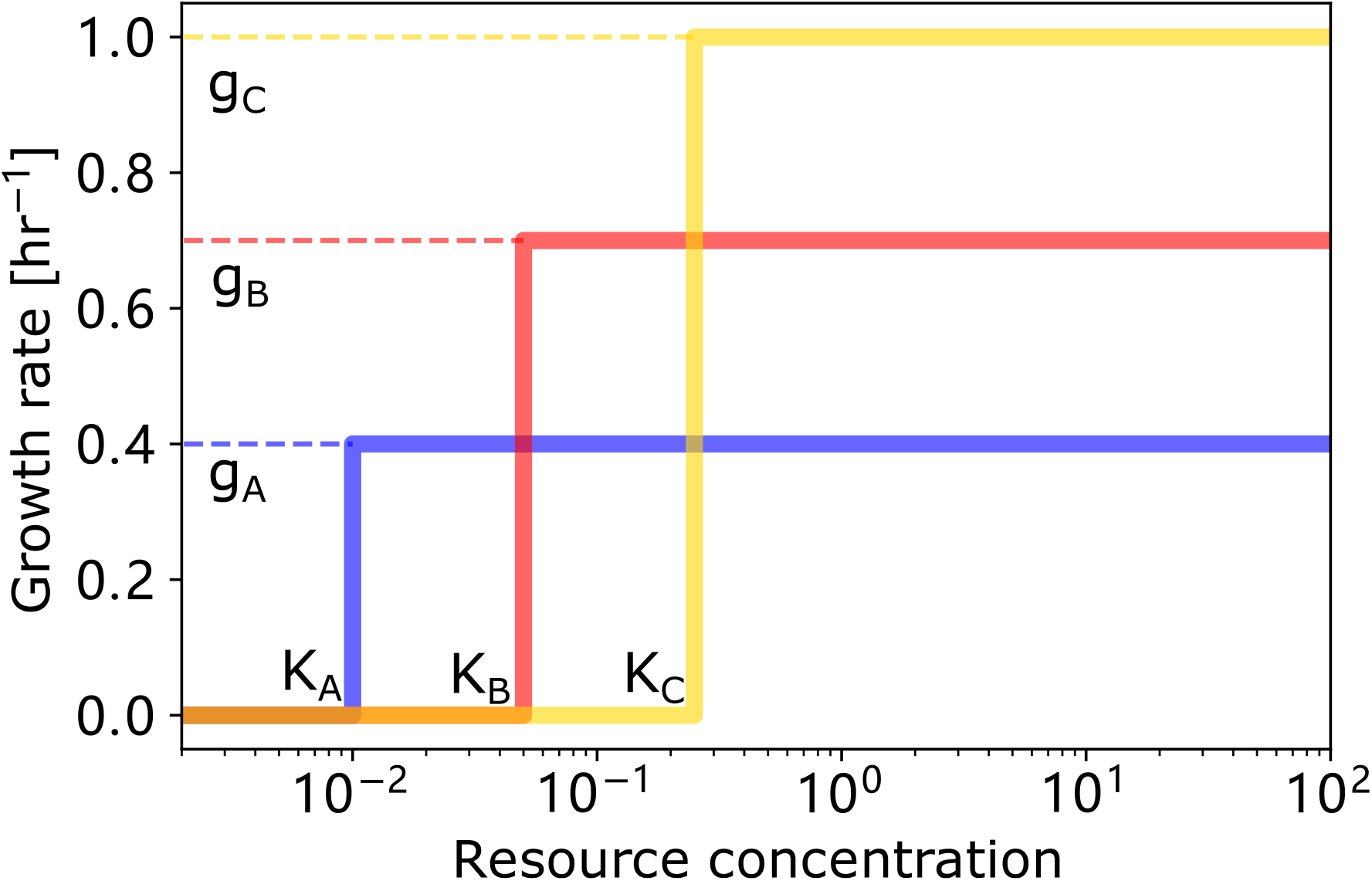
Growth curve of the simplified model. The 3 solid lines show how growth rate depends on the resource concentration in our simplified model. Here *g_A_*, *g_B_* and *g_c_* are approximations of the maximum growth rate from Monod’s law of growth, and *K_A_*, *K_B_* and *K_C_* are approximations of the substrate affinity.

*c*_0_ being the initial concentration of the resource at the beginning of a dilution cycle (obviously, it should be greater than *K*_1_).

Now *B*_1_ stops growing and *B*_2_ continues growing exponentially till the point *c* = *K*_2_. From that time point till the end of the dilution cycle the biomass conservation law reads simply as

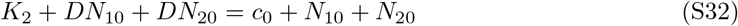

From (S32) we get, understandably, 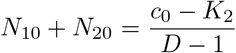, so that if coexistence indeed takes place, each of the steady state concentrations of both species should lie between 0 and 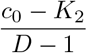. Using this consideration, from (S31) and (S32) we get

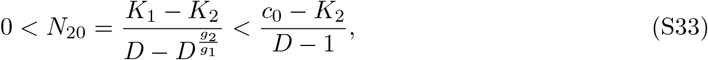

In the limit *c*_0_ ≫ *K*_2_ the coexistence condition takes the form

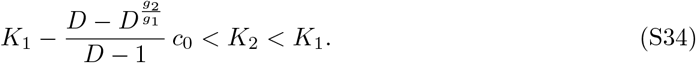

This is a fairy mild restriction, as a coexistence condition. For instance, if *g*_2_ → 0, the bacterium *B*_2_ will be able to invade for any *K*_2_ < *K*_1_ (though it may be present in a very small amount if *K*_2_ is close to *K*_1_). This is because at *c* = *K*_1_ the microbe *B*_1_ stops growing, and the biomass equal to *K*_1_ – *K*_2_ is always granted to *B*_2_. In fact, it is more restrictive for *B*_1_ that grows faster, but has larger *K*–constant. Real Monod’s dynamics does not work this way, and yet there are similarities, especially when the *K*–constants of both competitors are very different.

Suppose there are three microbes trying to share the same environment containing the single source c, under the same serial dilution conditions; for the sake of definiteness, we assume *g*_1_ > *g*_2_ > *g*_3_ and thus, to make coexistence possible, it should also happen that *K*_1_ > *K*_2_ > *K*_3_. If they succeed in sharing, then, at the steady state, the time point at which *B*_1_ stops growing must be *T*_1_ = log*D*/*g*_2_ (and at that time we should have *c* (*T*_1_) = *K*_1_; the microbe *B*_2_ stops growing must be *T*_2_ = log *D*/*g*_2_ (at *c*(*T*_2_) = *K*_2_) and, finally, *B*_3_ stops growing at the time point *T*_3_ = log *D*/*g*_3_ (at *c* (*T*_3_) = *K*_3_). Thus, instead of (S31) and (S32) we will have the following system of equations describing the biomass conservation at each of these three points:

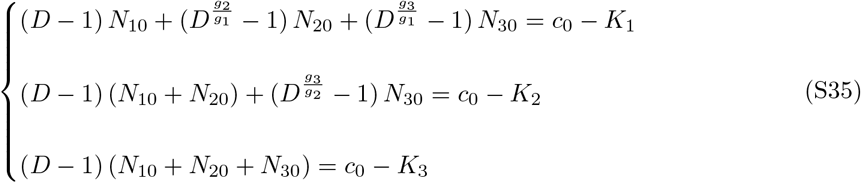

To begin with, it is obviously realizable, meaning that there exists a set of parameters (*g, K*), *c*, *D* that would ensure such coexistence. Take any set *g*_1_ > *g*_2_ > *g*_3_, *D* > 1, and for any positive numbers *N*_10_, *N*_20_, *N*_30_ the values *c*_0_ – *K*_1_, *c*_0_ – *K*_2_ and *c*_0_ – *K*_3_ will be obtained. Fix any *c*_0_ > (*D* – 1) (*N*_10_ + *N*_20_ + *N*_30_) and get *K*_1_, *K*_2_, *K*_3_ < *c*_0_ ordered as required. Next, in this model, it is even quite easily realizable. Solving the system (S35) we get for *B*_3_

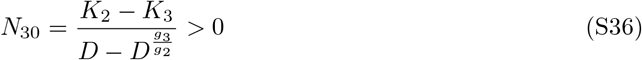

then for *B*_2_

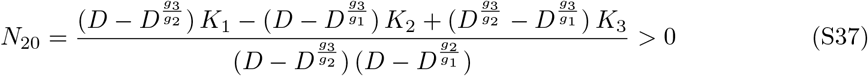

and, finally, for *B*_3_

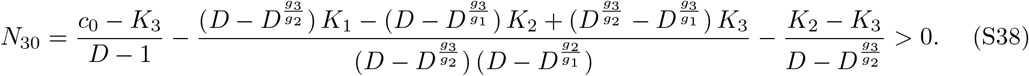

It can be seen from above that the conditions are still fairly easy to satisfy if *c*_0_, *K*_1_, *K*_2_ and *K*_3_ are well separated.

Coexistence of *n* microbes *B*_1_, *B*_2_,…, *B_n_* is governed by the system of equations

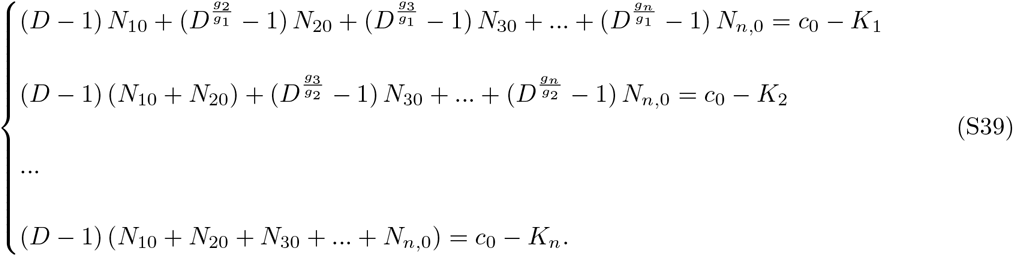

As with (S35), it is easy to see that such a coexistence is realizable. Take any set of the parameters *g*_1_,…, *g_n_*, any *D* > 1 and any set of positive *N*_10_,…, *N_n,0_*, assume any *c*_0_ > (*D* – 1) (*N*_10_ + *N*_20_ + *N*_30_ +… + *N*_n,0_) and get *K*_1_,…, *K_n_*, again automatically ordered as required.

In this model, any new survivor comes in relatively easily, and indeed we get a large number of microbes surviving on a single source in our simulations. It is not so with the Monod’s dynamics, with which it is hard to get two species coexisting, much harder yet to get three and almost impossible further on. Nevertheless, in principle, the parameters can be found to get such a coexistence. To see why it should happen, break the Monod’s curve into a sequence of flat Heaviside’s steps, so that it is approximated by a “stairs”, each of the steps corresponding to a new value of the

**Figure S4:**
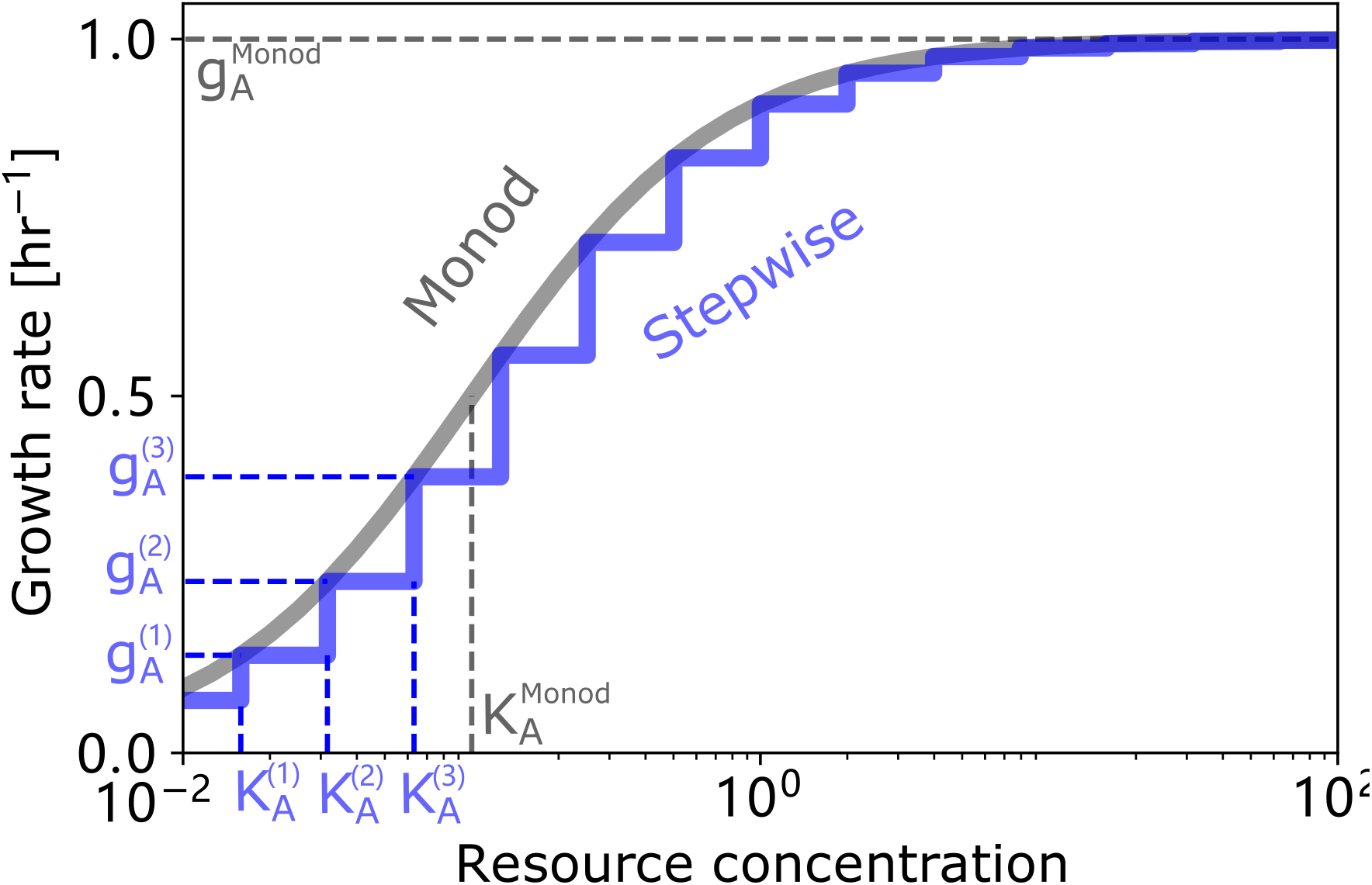
Monod growth curve approximated by the simplified model. The grey line shows the growth curve of a strain that obeys Monod’s law of growth on a single resource, with maximum growth rate 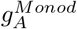 and affinity 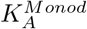. The blue line is the growth growth curve of a strain that grows on a set of unique resources and follows our simplified growth model with growth rates 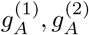… etc. and affinity 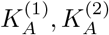… etc.

logarithmic growth rate. In this way, we look at the single resource c as a set of a number of unique resources, each providing its own value of the (exponential) growth rate for each microbe. In the study, ref. [25] we show that, in principle, in a serial dilution setting it is possible in this case to have the number of survivors matching the number of sources. However, it works best when the microbes have different source preferences: say, one of them starts consuming the resource number 2 first, while the other one starts with the resource number 1 etc. In our case, the species are bound to have the same food preferences, and in this case a multiple coexistence is much harder (though possible) to arrange.

Note that, even if breaking the Monod’s curve in any number of steps we could hope for any number of survivors, we play on a shaky ground. As the range of the parameters favoring the coexistence of more and more species becomes smaller, we can well fall out of the accuracy limits of the Monod’s dynamics. Being in itself an approximation, it cannot support that sort of handling to the infinity.

## Notes

https://github.com/maslov-group/Coexistence_of_g_and_K

### Competing Interest Statement

The authors have declared no competing interest.

### Summary of Updates

Revised references.

